## Supplementary Figures for "Structural Insights into the Dynamics of Water in SOD1 Catalysis and Drug Interactions"

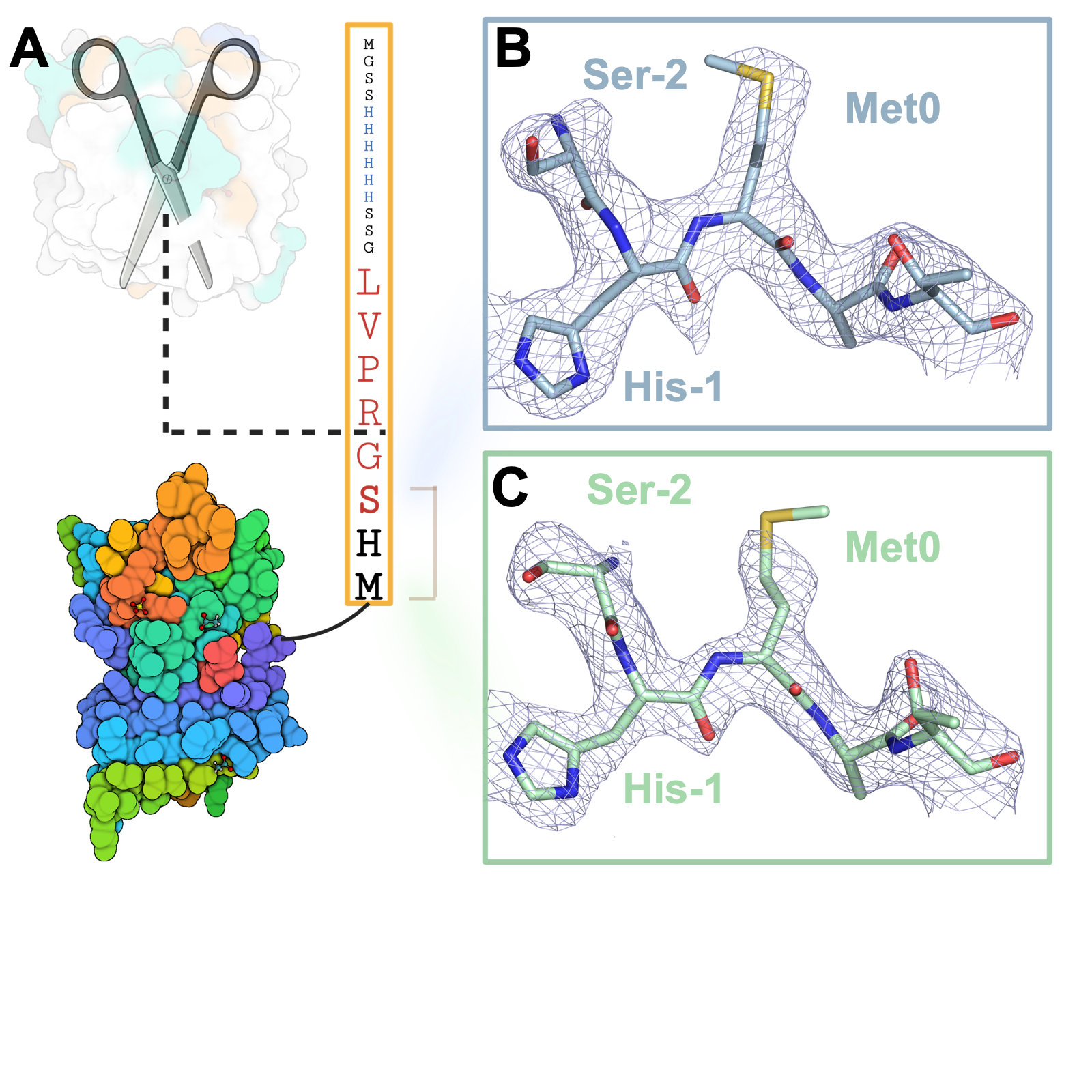

**Supplementary Figure 1. Thrombin-specific N-terminal cleavage of hSOD1. A)** The modified N-terminal of hSOD1 is cleaved by thrombin, leaving 3 residues in chain B, in blue **(B)** and D, in mint **(C)**. The *2Fo-Fc* electron density map of the remaining residues is contoured at 1σ level and colored in dark blue.

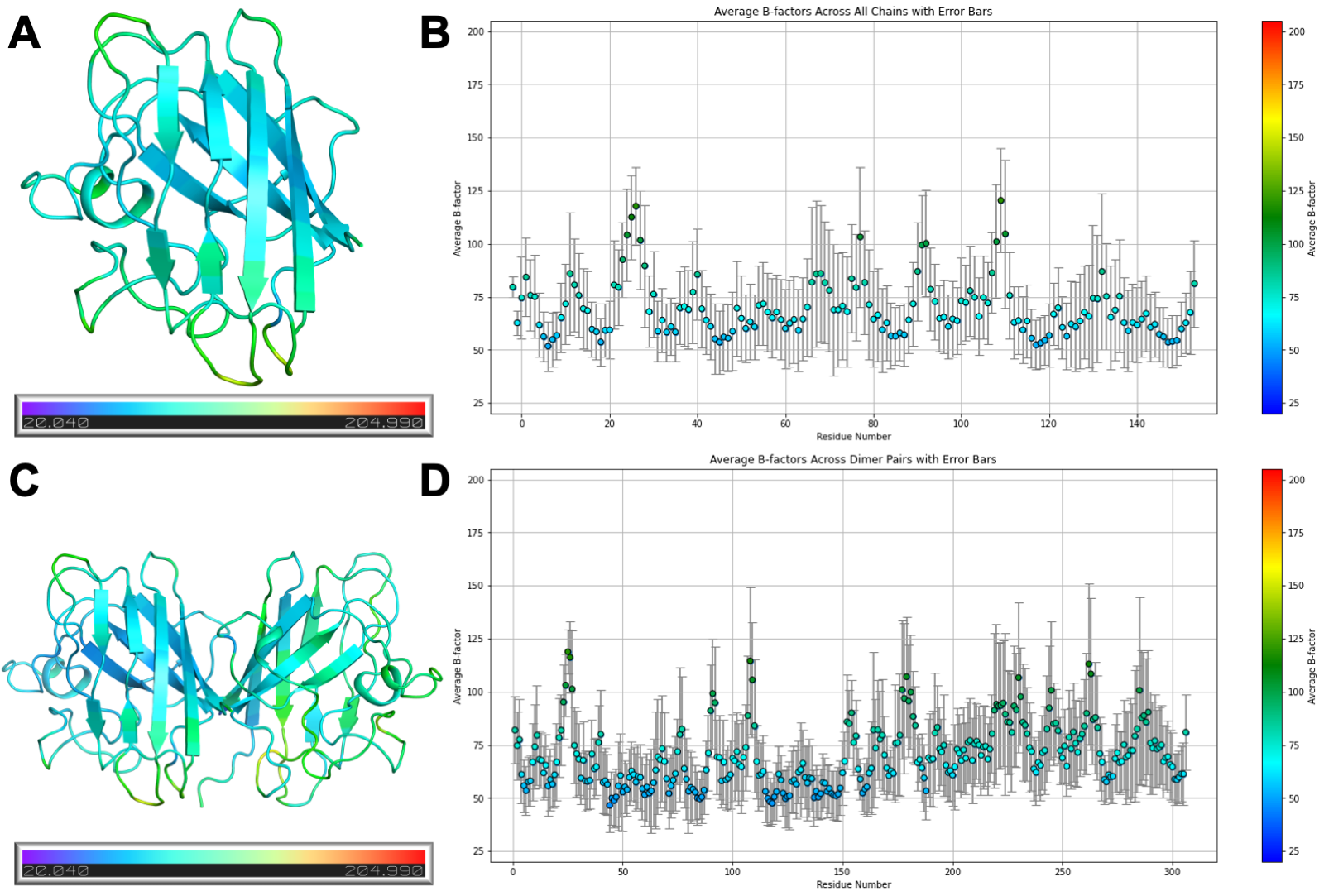

**Supplementary Figure 2. The average *B*-factors of the monomeric and dimeric forms of hSOD1.** The average *B*-factor per residue is calculated to be mapped on the structure and each value is displayed on a scatter graph for monomeric **(A, B)** and dimeric (**C, D)** forms of the hSOD1. In panel **D**, the first 153 residues correspond to the first monomers (A, C, E, G, I, K chains), while the second 153 residues correspond to the second monomers (B, D, F, H, J, L chains). The minimum value is 20.040 in blue, while the maximum value is 204.990, as indicated by red in the spectrum, for consistency.

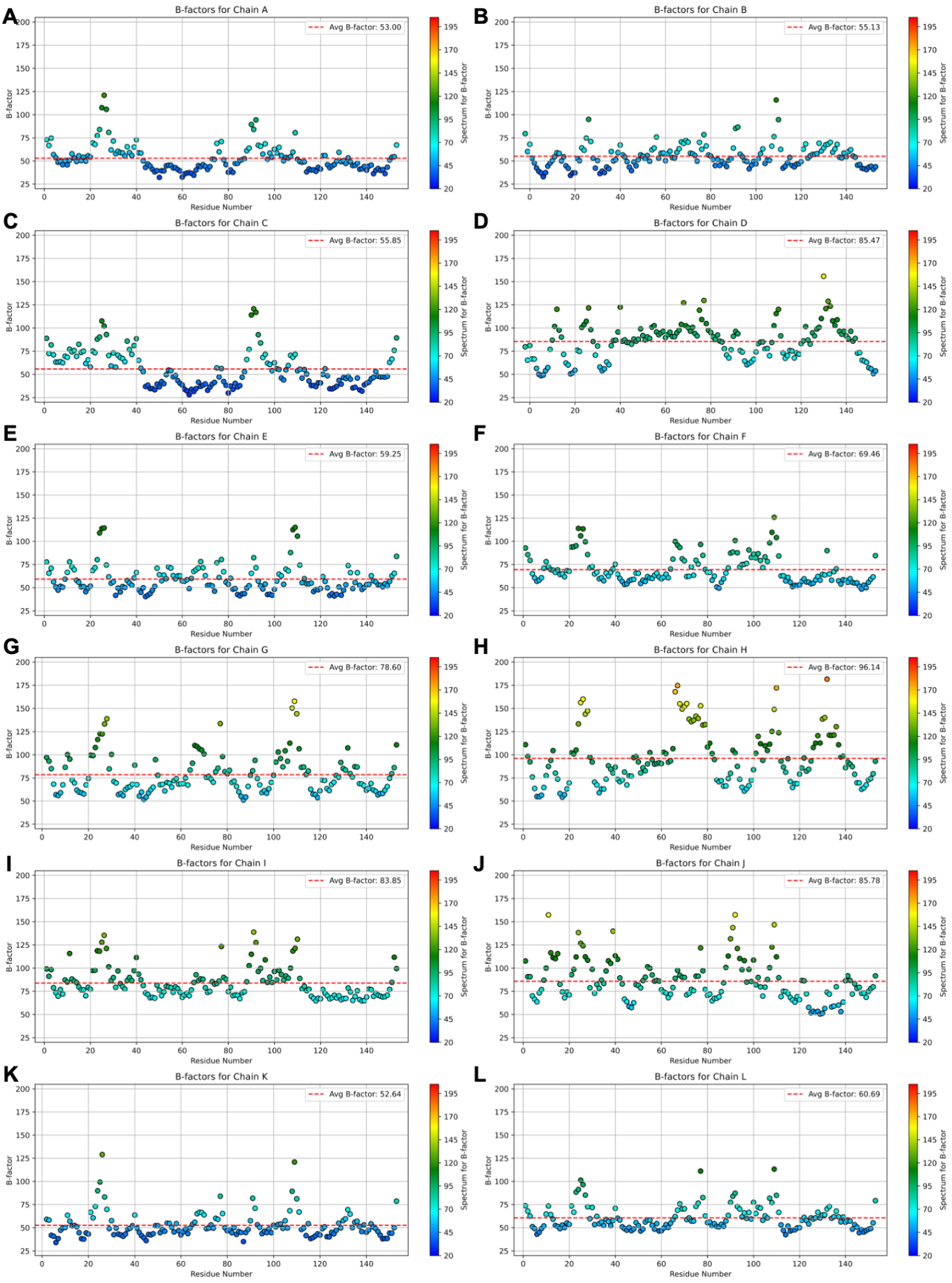

**Supplementary Figure 3. Scatter plot of all-atom *B*-factors for each residue across twelve different chains.** All residues in each chain are colored according to the consistent spectrum of a minimum value of 20.040 with blue color, and a maximum value of 240.990 with red color. The dashed line in each graph represents the average B-factor, calculated specifically for each chain.

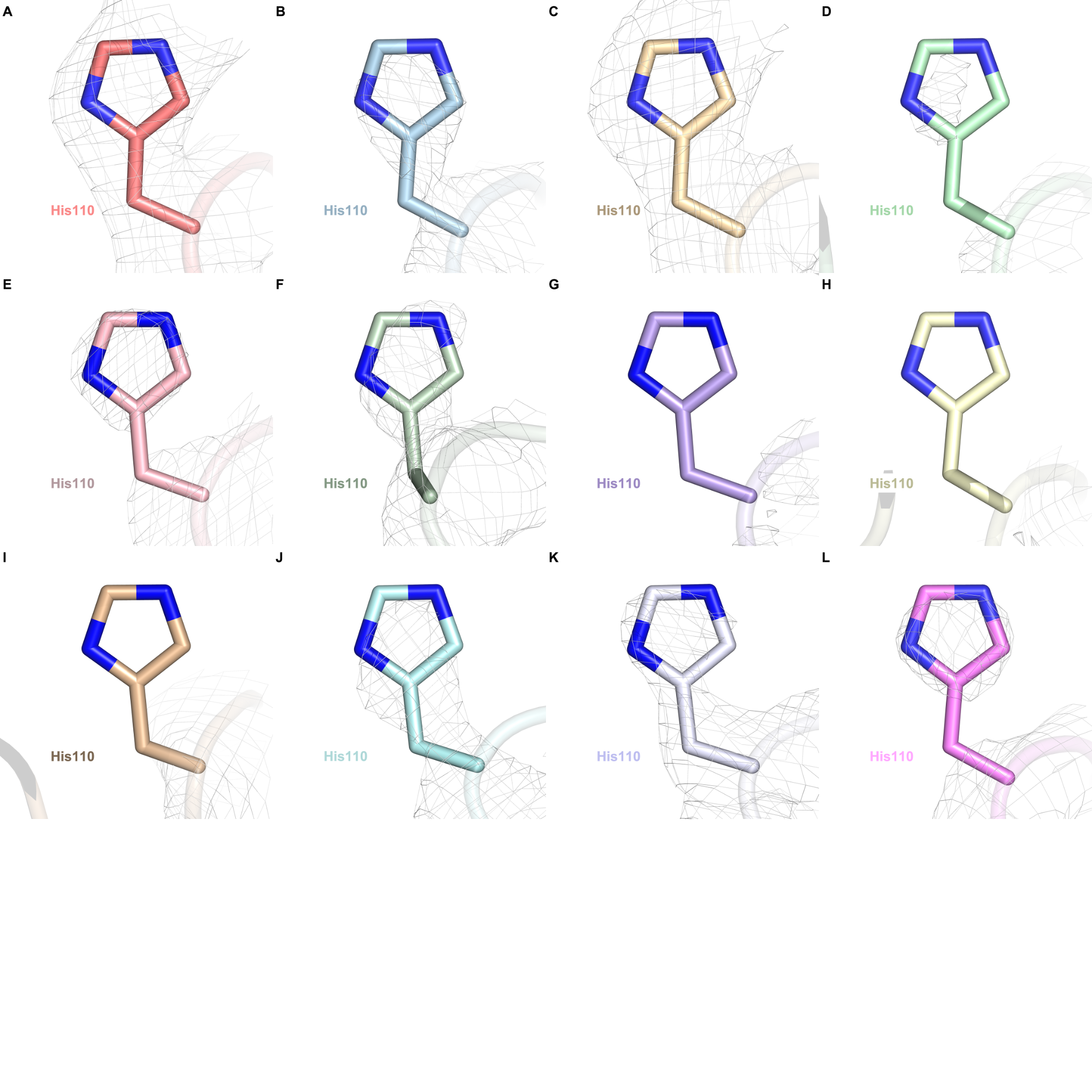

**Supplementary Figure 4. The *2Fo-Fc* electron density map for the flexible His110 residue.** The all-features map of His110 residues from each chain is contoured at 1σ level and colored in gray. His110 residues coordinated by Zn^CRYST^ **(A,C)** have a better-defined electron density compared to free histidines.

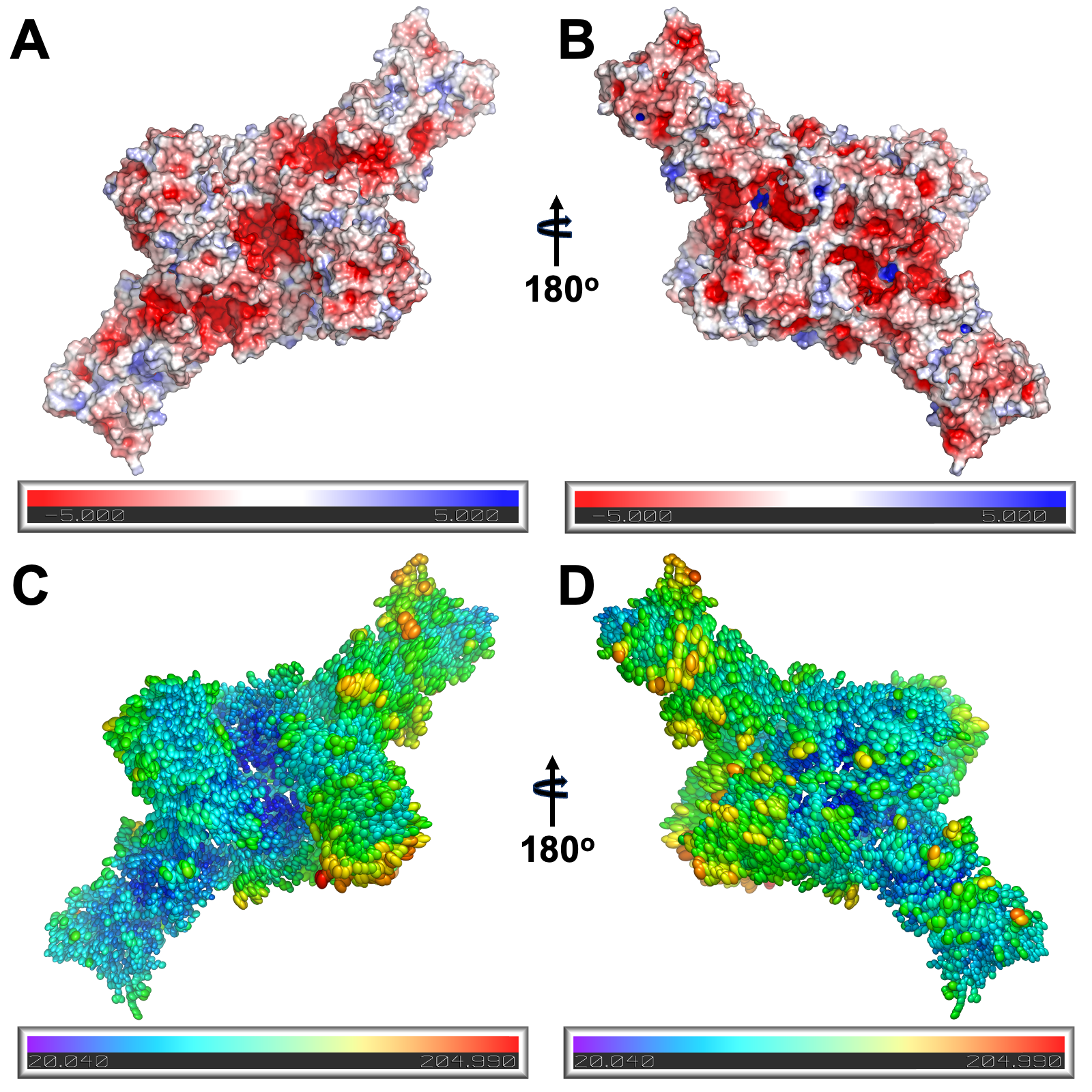

**Supplementary Figure 5. Representation of electrostatic surface potential and ellipsoids of the hSOD1 structure. A,B)** APBS electrostatics in *PyMOL* is used to detect the highly acidic core and **C,D)** anisotropic *B*-factors for all atoms are represented by thermal ellipsoids revealing the rigid core of the crystal structure.
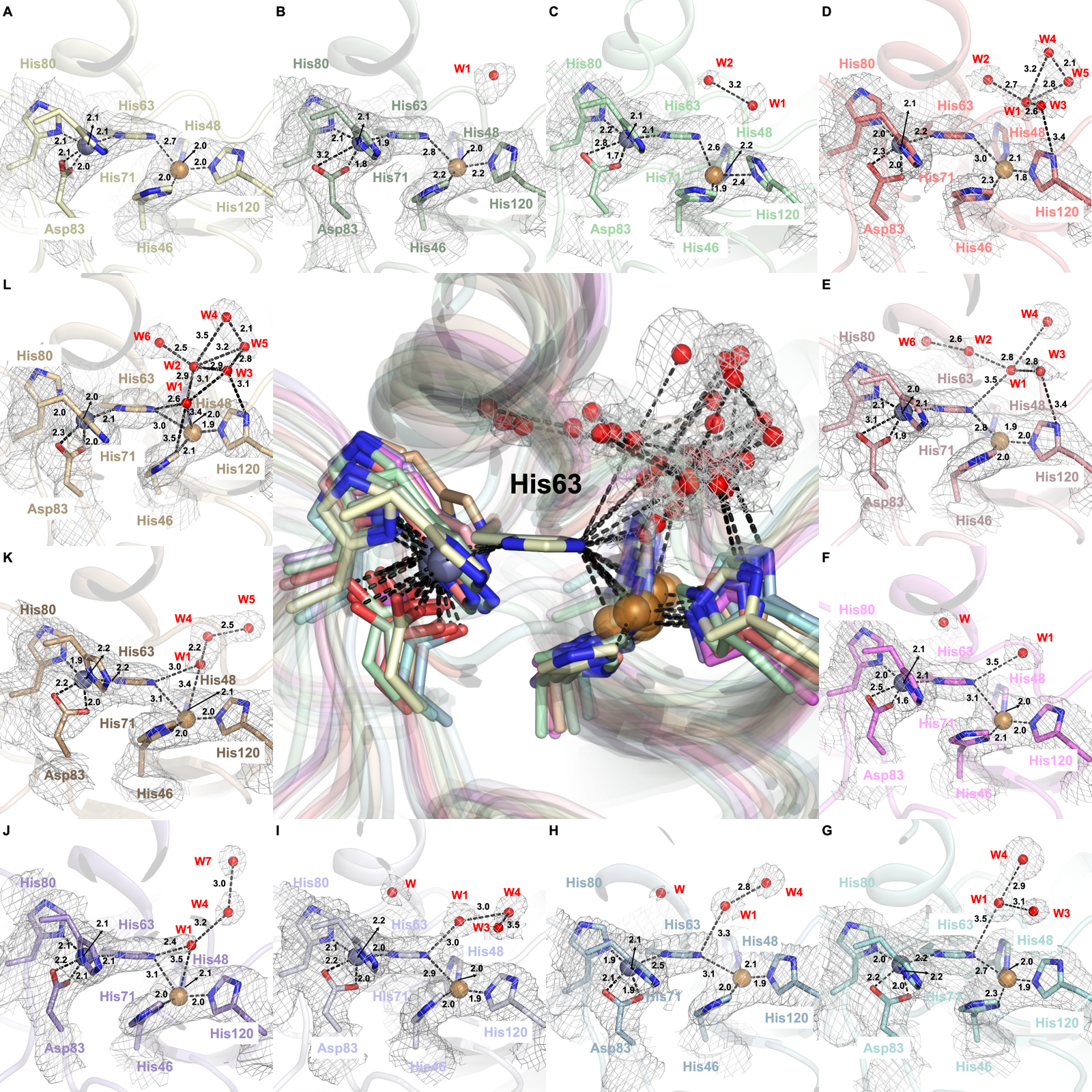

**Supplementary Figure 6. The water coordination and bond lengths at twelve hSOD1 active sites.** The center panel shows the superpositions of all 12 active site regions, each centered on the CE1 atom of His63. It reveals the positional shifts of active site water molecule dynamics, the distances between the residues are given in angstroms (Å). **A-K)** panels are ordered to show the choreography of the water molecules approaching and departing from the active site during the catalytic cycle. The *2Fo-Fc* electron density map covering waters and other active site residues is contoured at 1σ level and colored in gray.

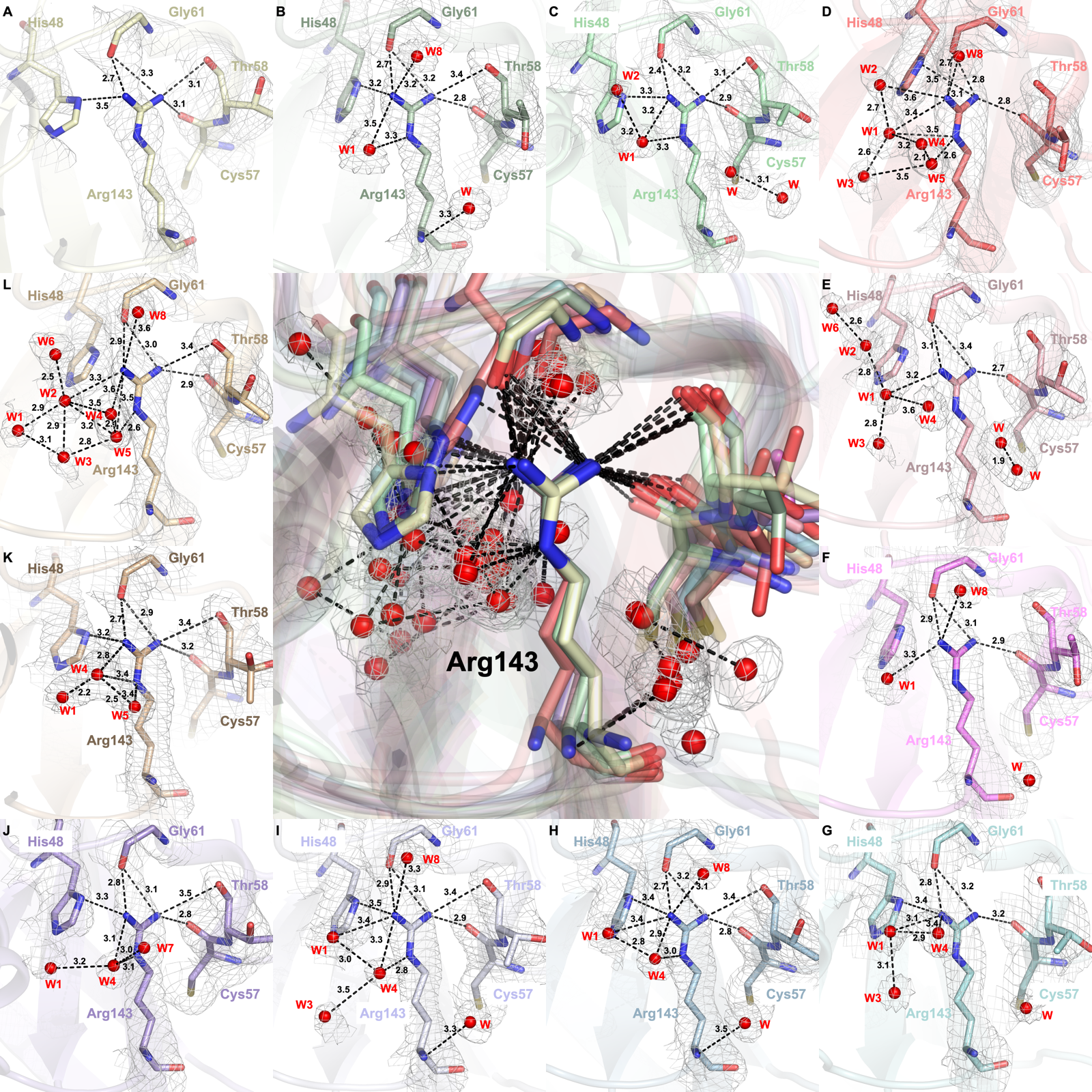

**Supplementary Figure 7: Allosteric water coordination and bond lengths around the Arg143 residue.** The center panel shows the superpositions of all 12 active site regions, each centered on the NE atom of Arg143. **A-K)** panels are ordered to show the choreography of the water molecules approaching and departing from the active site during the catalytic cycle. The *2Fo-Fc* electron density map covering waters is contoured at 1σ level and colored in gray.
